# Fusion of a non-specific DNA-binding domain enhances Cas12a trans-cleavage for robust nucleic-acid diagnostics

**DOI:** 10.64898/2026.05.30.729025

**Authors:** Diogo Figueiredo, Mitsuru Hattori, Tetsuichi Wazawa, Martijn Zwama, Kunihiko Nishino, Takeharu Nagai

## Abstract

CRISPR–Cas12a underlies powerful genome-editing and nucleic-acid detection technologies, yet its performance is limited by inefficient target engagement and low catalytic turnover, particularly at low target abundance and elevated temperatures. Here, we report a modular protein-engineering strategy to enhance Cas12a trans-activity by fusing the hyperthermophilic DNA-binding protein Sso7d to the N terminus of *Lachnospiraceae bacterium ND2006* Cas12a (LbaCas12a). The resulting fusion enzyme shows a twofold improvement in detection sensitivity, a 4.6-fold increase in *k*_cat_, and substantially accelerated trans-cleavage kinetics compared with wild-type Cas12a. These enhancements are observed across multiple guide RNAs, diverse DNA substrates, and a broad temperature range from 37 to 60 °C. The engineered Cas12a enables robust detection of the intrinsic *blaOXA-51* and acquired *blaOXA-24* antibiotic resistance genes from multiple *Acinetobacter baumannii* strains with independently validated resistance profiles, whereas wild-type Cas12a produces little or no detectable signal. Signal output is increased by up to ∼30-fold, expanding the effective detection window and shortening time-to-positive. Together, these results establish fusion of an accessory DNA-binding domain as a proof-of-concept for Sso7d fusion to LbCas12a and that generalization to other CRISPR effectors or non-specific DNA-binding domains represents an exciting avenue for future investigation.

## INTRODUCTION

CRISPR–Cas systems have transformed molecular diagnostics by enabling programmable nucleic acid recognition coupled to collateral nuclease activities that generate amplified, easily detectable signals ^1,2^. Among these platforms, *Lachnospiraceae* bacterium ND2006 Cas12a (Cpf1) has emerged as a particularly attractive enzyme for DNA detection owing to its robust trans-cleavage activity and minimal protospacer-adjacent motif (PAM) requirements ^3^. Beyond diagnostics, Cas12a is a versatile RNA-guided endonuclease widely used for genome editing ^4^, transcriptional regulation ^5,6^ and molecular sensing ^7^, underscoring its broad utility across basic and applied biology.

Upon target recognition, Cas12a undergoes a large conformational rearrangement that activates its RuvC nuclease domain ^8^, enabling both sequence-specific cis-cleavage of double-stranded DNA and promiscuous trans-cleavage of single-stranded DNA reporters ^9,10^. This target-activated trans-cleavage underpins widely used diagnostic platforms such as DETECTR ^1,7,11^ and related isothermal amplification workflows. Despite these strengths, Cas12a performance is frequently limited under practical assay conditions. In particular, signal generation becomes inefficient at low target abundance or elevated temperatures—regimes that are central to rapid, high-sensitivity diagnostics.

Multiple factors contribute to these limitations. Efficient Cas12a activation requires stable crRNA– target duplex formation ^12,13^ yet many guide–target combinations exhibit slow binding kinetics or incomplete activation ^14^. In addition, reduced target engagement and partial thermal destabilization above physiological temperatures constrain compatibility with high-temperature amplification methods such as loop-mediated isothermal amplification (LAMP) or recombinase polymerase amplification (RPA) workflows ^15–17^. Finally, the intrinsic catalytic turnover of Cas12a (*k*_cat_) is modest, limiting time-to-result and sensitivity in applications demanding rapid signal amplification. Strategies that enhance Cas12a catalysis while preserving specificity, therefore, remain a central challenge for CRISPR-based diagnostics.

Most reported approaches to improve Cas12a performance rely on guide RNA engineering ^18,19^, optimization of targets or reporter substrates ^20^, or the use of chemical additives ^21^. Protein engineering strategies have largely focused on random mutagenesis, directed evolution ^22,23^, or rational redesign of active-site residues ^24,25^ — approaches that are often labor-intensive and difficult to generalize. By contrast, comparatively little attention has been given to modular domain fusion as a means to enhance CRISPR enzyme performance, despite emerging evidence that accessory domains can tune target engagement or stabilize activated conformations ^26^.

The hyperthermophilic DNA-binding protein Sso7d is a small (7 kDa), basic, and exceptionally stable protein that binds double-stranded DNA with minimal sequence preference by positioning its triple-stranded β-sheet across the minor groove ^27,28^. Sso7d remains folded at extreme temperatures (T_m_ = 100.2 °C) ^29^ and across a broad pH range (pH 0–12) ^30^, making it an attractive modular element for engineering DNA-processing enzymes. Notably, Sso7d binds DNA with nanomolar affinity under salt conditions compatible with maximal Cas12 activity ^31,32^. Whether tethering such a nonspecific DNA-binding domain to Cas12a could enhance target engagement, catalytic turnover, or thermal robustness, however, has not been explored in detail ^26^

Here, we show that fusion of the nonspecific DNA-binding domain Sso7d to Cas12a yields a highly active Cas12a variant with markedly enhanced analytical performance. This engineered fusion enables up to a 30-fold increase in detectable trans-cleavage signal during the detection of antimicrobial resistance (AMR) genes from clinical *Acinetobacter baumannii* isolates. The improved activity is maintained across multiple guide RNAs and substrate types, accelerates time-to-positive, and enhances detection sensitivity without compromising target specificity. Together, these results demonstrate that DNA-binding domain fusion provides a proof-of-concept to expand the utility of Cas12a for sensitive and rapid molecular diagnostics.

## RESULTS

### Engineering and functional screening of Sso7d–Cas12a fusion variants

To test whether accessory DNA-binding domains can potentiate Cas12a activation, we engineered a systematic panel of Sso7d–Cas12a fusion variants encompassing N-terminal and C-terminal fusion topologies, split-Cas12a architectures, and multiple linker designs (**Fig. 1**). For N-terminal fusions (Sso7d-Cas12a in **Fig. 1**), the DNA-binding domain was connected to Cas12a using an extended XTEN linker (an artificial extended and unstructured linker that has seen broad utility for creating chimeric fusions with Cas proteins with the following sequence – SGSETPGTSESATPES)^33–35^ to provide conformational flexibility and minimize steric interference with RNP assembly. For C-terminal fusions (Cas12a-Sso7d in **Fig. 1**), both short flexible glycine–serine (GGS) linkers and extended XTEN linkers were evaluated. An N-terminal maltose-binding protein (MBP) tag was included in selected constructs to enhance soluble expression. Reports have previously shown that the MBP tag has no effect on the cleavage efficiency of LbaCas12a (^36^). For split Sso7d fusion (Sso7d fusion(N) + Sso7d fusion in **Fig. 1**), N-terminal fusions used an XTEN linker and the Cas12a protein was split at site 575. All constructs were heterologous expressed in *Escherichia coli (***Supplementary Fig. 1 and 2**) and subjected to an initial functional screen based on target-activated trans-cleavage activity using a fluorogenic single-stranded DNA reporter, providing a rapid and quantitative readout of Cas12a activation prior to detailed kinetic characterization.

**Figure 1.**
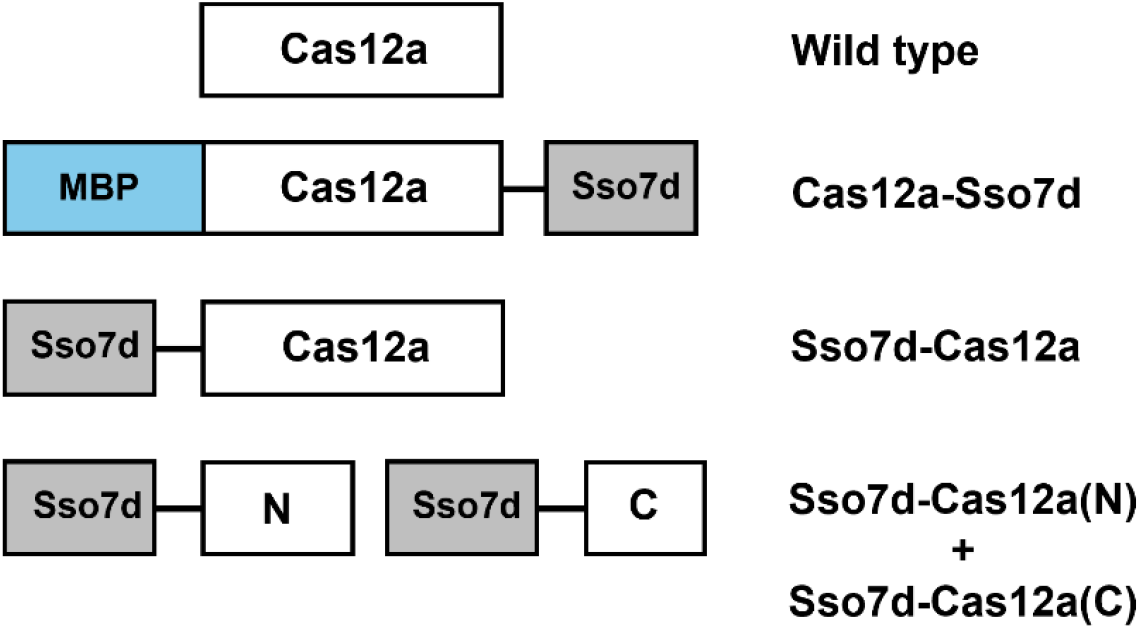
Design and architecture of engineered Cas12a fusion proteins. Schematic representation of Cas12a fusion constructs illustrating the fusion sites, orientations, and linker designs of the DNA-binding protein relative to Cas12a. N-terminal fusions were generated using an XTEN linker, whereas C-terminal fusions were constructed with either GGS or XTEN linkers and included an MBP tag to enhance solubility. Schematics depict construct architecture only and are not drawn to scale.

To enable consistent benchmarking across constructs, guide RNAs were designed to target the ampicillin resistance gene (AmpR), present in plasmids such as pRSET_B_. A PCR-amplified fragment encompassing this locus was used as the activating double-stranded DNA target in all subsequent assays. Among the diverse fusion architectures evaluated, a single configuration—fusion of Sso7d to the N terminus of Cas12a via an extended XTEN linker—exhibited a pronounced enhancement in target-activated trans-cleavage activity, under conditions of saturating target DNA (300 nM DNA, 30 nM Cas12a Ribonucleoprotein complex) (**Fig. 2a**). The optimized fusion construct displayed up to a fourfold increase in initial reaction velocity compared to wild-type Cas12a (**Fig. 2b**). Notably, this enhancement was observed without any detectable increase in background fluorescence or target-independent activation, indicating that fusion of Sso7d selectively enhances productive target engagement and activation rather than destabilizing the inactive enzyme state. Based on its superior performance and favourable signal-to-background characteristics, this construct—hereafter referred to as Sso7d–Cas12a—was selected for comprehensive kinetic, specificity, and limit-of-detection analyses.

**Figure 2.**
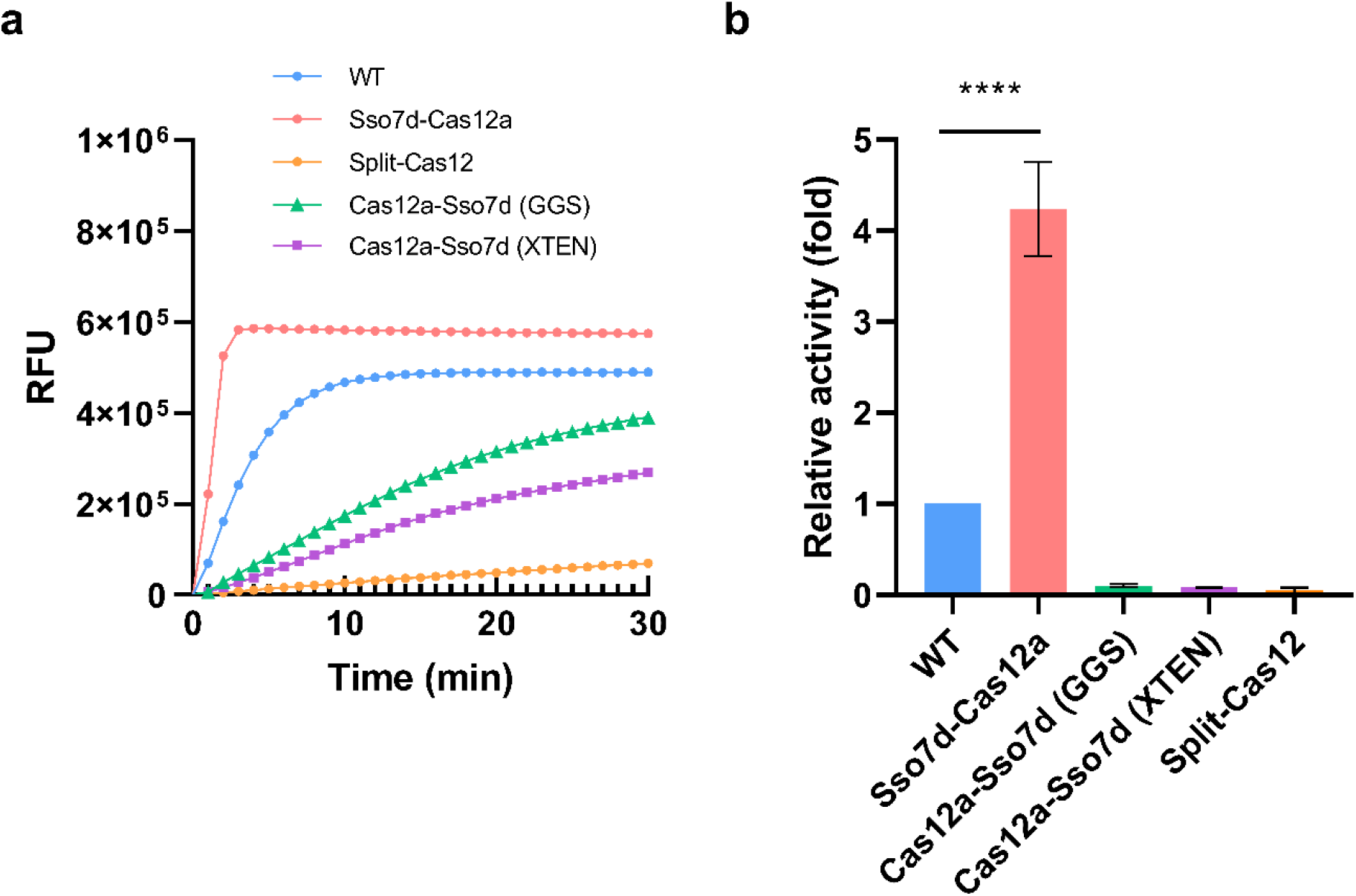
Comparative activity and kinetic behaviour of engineered Cas12a fusion constructs. (a) Representative kinetic traces showing time-dependent DNA cleavage for the corresponding constructs (background subtracted). (b) Trans-cleavage activity of wild-type Cas12a and engineered Cas12a fusion constructs measured under identical reaction conditions. Activities were calculated from initial reaction velocities (at R^2^=0.99) and normalized to wild-type Cas12. RFU – Relative fluorescent units; WT – Wild type; Reactions were performed at 37 °C for 1 h, with fluorescence recorded every 1 min. Data represent mean ± s.d. from at least three independent experiments.

### Sso7d fusion enhances catalytic turnover and thermal robustness

We next quantified the effect of Sso7d fusion on Cas12a catalysis by performing steady-state kinetic analyses at 37 °C and 60 °C using an excess fluorogenic single-stranded DNA reporter (**Fig. 3a and 3c**, respectively). Reaction velocities were extracted from the linear phase of the fluorescence time courses (R^2^=0.99) across a range of different DNA target concentrations (300 nM to 0.06 nM) and fitted to the Michaelis–Menten equation to obtain enzymatic parameters (**Table 1**). The respective values of ΔRFU/min in this region were used to determine and calculate activities (relative to the wild type – WT). At 37 °C, the Sso7d–Cas12a fusion exhibited a 4.6-fold increase in *k*_cat_ relative to wild-type Cas12a, while the apparent *K*_m_ was only modestly affected (1.46-fold increase). This indicates that the main effect of Sso7d fusion is on the catalytic step rather than on substrate binding. If the fusion protein acted by increasing DNA binding affinity, a decrease in *K*_m_ would be expected. The small *K*_m_ increase is instead consistent with a minor effect on RNP–target assembly, while the large *k*_cat_ increase points to faster cleavage once the complex is formed. This pattern suggests that the rate-limiting step in the wild-type reaction is the conformational change that activates the RuvC domain after target binding, and that this step is accelerated by Sso7d fusion. t_sat_, the time at which each fluorescence trace reached 95% of its fitted asymptotic maximum, was 3 minutes for Sso7d–Cas12a compared to 15 minutes for wild-type Cas12a, confirming that the fusion enzyme reaches signal plateau substantially faster under identical reaction conditions (**Table 1**). A similar trend was observed at 60 °C, where the fusion construct retained elevated catalytic rates compared to wild-type Cas12a (**Fig. 3b, 3d**), demonstrating that the activity enhancement persists under elevated temperature conditions. The cis cleavage activity of the Sso7d-Cas12 protein was also evaluated (**Supplementary Fig. 3**). At both temperatures (37 ºC and 60 ºC), cleavage of the target DNA (linear dsDNA) was higher when compared to the wild type (**Supplementary Fig. 3**), with an approximately 1.5-fold improvement at 37°C. This is smaller than the 4.6-fold improvement in trans-cleavage *k*_cat_ measured under the same conditions. Both cis- and transcleavage require R-loop formation but differ in the subsequent cleavage step: cis-cleavage involves cutting the double-stranded target DNA, while trans-cleavage acts on the single-stranded reporter. The larger improvement in trans-cleavage compared to cis-cleavage suggests that Sso7d has an effect beyond R-loop formation, specifically on RuvC cleavage of single-stranded DNA substrates. These results show that fusion of the accessory DNA-binding domain preserves dsDNA cis-cleavage activity in the fusion construct while improving the catalytic efficiency of Cas12a across a broad temperature range.

**Table 1.** Kinetic parameters of wild-type Cas12a and Sso7d–Cas12a.

| Protein | $K_{cat}$ ($\text{min}^{-1}$) | $K_m$ (nM) | $K_{cat}/K_m$<br>($\text{min}^{-1} \text{nM}^{-1}$) | $t_{sat}$ (min) |
| --- | --- | --- | --- | --- |
| Sso7d-Cas12a | 38.3 | 117.8 | 0.33 | 3 |
| WT | 8.3 | 80.59 | 0.1 | 15 |

**Figure 3.**
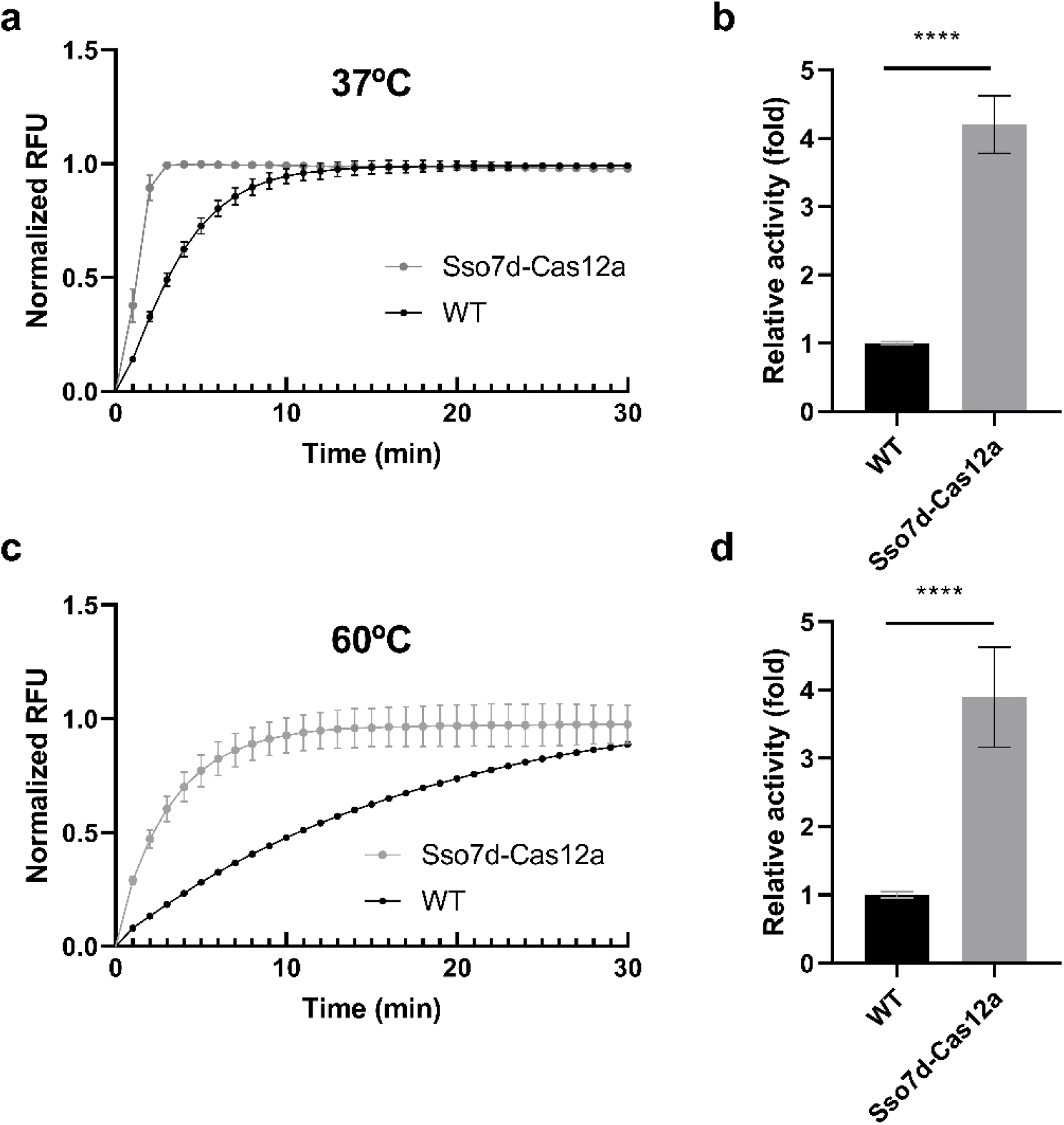
Temperature-dependent enhancement of Cas12a catalytic activity by DNA-binding domain fusion. (a) Steady-state trans-cleavage kinetics of wild-type Cas12a and Sso7d-Cas12a at 37 °C under multiple-turnover conditions with excess substrate DNA. (b) Corresponding relative trans-cleavage activity of Cas12a fusion variants normalized to wild-type Cas12a (ΔRFU/min relative to WT). (c) Steady-state trans-cleavage kinetics at 60 °C. (d) Corresponding relative trans-cleavage activity of Sso7d-Cas12a normalized to wild-type Cas12a (ΔRFU/min relative to WT). WT – Wild type; RFU – Relative fluorescent Units; Reactions were monitored for 1 h with fluorescence recorded every 1 min. Data represent mean ± s.d. from at least three independent replicates.

### Catalytic enhancement varies with guide RNA and is substrate-independent

To assess whether the observed enhancement depended on guide RNA sequence, we evaluated a diverse panel of crRNAs (**Supplementary Table 1**) targeting different protospacer sequences (TTTA, TTTG) within the same double-stranded DNA target (**Supplementary Table 3**). Sso7d–Cas12a showed significantly higher catalytic rates than wild-type Cas12a for guide RNAs 2 and 3, while guide RNAs 1 and 4 showed no statistically significant improvement (**Fig. 4a**). The degree of enhancement varied across guides and appeared to correlate with baseline activity: guide RNAs with lower activity against wild-type Cas12a showed larger improvements with the fusion protein. This suggests that Sso7d is most effective when the baseline rate of RNP–target activation is low, which has practical implications for guide RNA selection in diagnostic assays. We next evaluated the performance of Sso7d–Cas12a across distinct substrate architectures using the best performing guide RNA (guide RNA 2), including linear single-stranded DNA reporters and circular double-stranded DNA targets. Notably, increased activity was reflected both in relative cleavage levels and in accelerated reaction rates, indicating that the accessory DNA-binding domain enhances productive Cas12a activation across structurally distinct nucleic acid substrates (**Fig. 4b**). Across both substrate types, Sso7d-Cas12a consistently exhibited enhanced activation relative to wild-type Cas12a, demonstrating that the observed activity gain is largely independent of substrate topology. Even though Sso7d binds primarily dsDNA, some reports have shown that weak interactions with ssDNA can also occur under specific conditions ^37^

**Figure 4.**
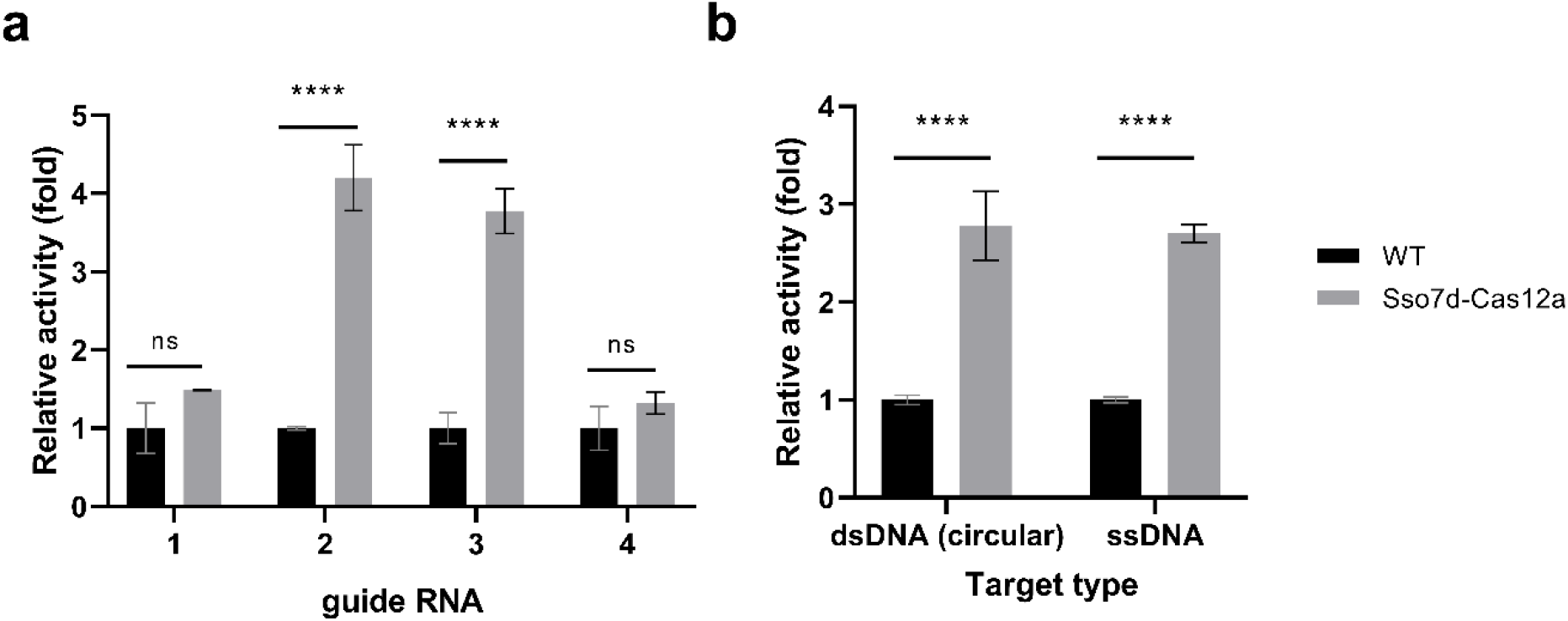
On-target and substrate dependent trans-cleavage activity of Cas12a fusion protein. (a) Relative trans-cleavage activity of wild-type Cas12a and Sso7d–Cas12a measured using on-target DNA substrate for each guide RNA. (b) Relative trans-cleavage activity of wild-type Cas12a and Sso7d–Cas12a measured using circular double-stranded DNA (dsDNA, AmpR) and single-stranded DNA (ssDNA) substrates for guide RNA 2. Reactions were performed at 37 °C for 1 h, with fluorescence recorded every 1 min. Bars represent mean ± s.d. from at least three independent replicates.

Specificity was further evaluated using another commonly used antibiotic resistance gene (Kanamycin resistance gene, KanR, OFF target). Under identical reaction conditions, Sso7d–Cas12a maintained target discrimination and improved specificity across all guide RNAs relative to wild-type Cas12a (**Fig. 5**). For the wild-type Cas12a, a significant increase in trans-cleavage RFU was observed under off-target conditions relative to NTC for guide RNAs 1 and 2 (**Fig. 5a, 5b**), whereas no significant difference between NTC and off-target conditions was detected for any guide RNA when using Sso7d-Cas12a. These results indicate that fusion of the accessory DNA-binding domain enhances catalytic efficiency without compromising, and improving, target specificity.

**Figure 5.**
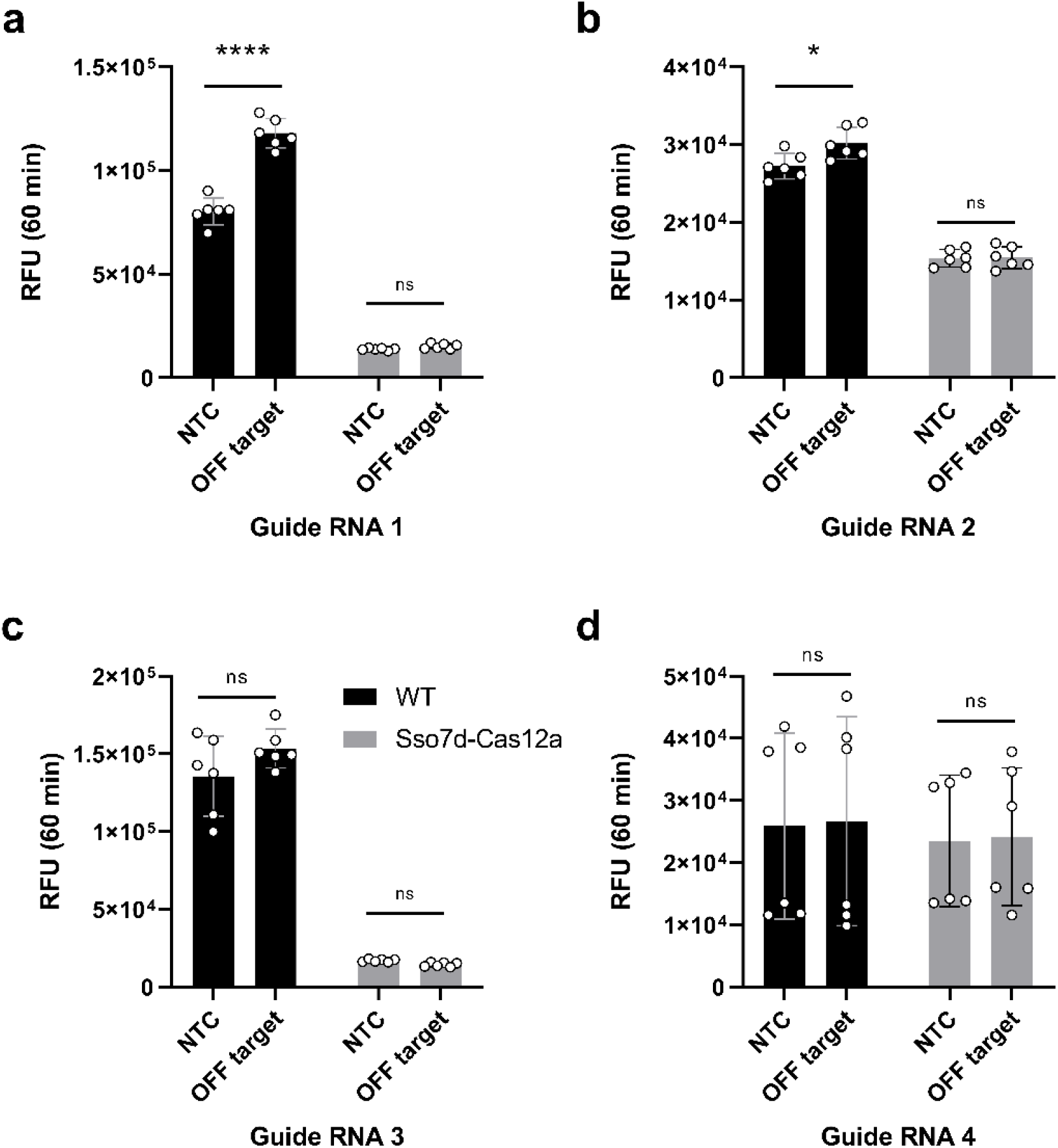
Guide RNA–dependent off-target activity of WT and fusion Cas12a. Off-target signals were measured for four independent guide RNAs (a–d) in the presence (OFF-target) and absence (NTC) of the KanR dsDNA amplicon using WT or fusion Cas12a. NTC – Non-template control; OFF target – Kanamycin resistance gene; Reactions were performed at 37 °C for 1 h, with fluorescence recorded every 1 min. Data represent mean ± s.d. from six independent replicates.

To further examine whether increased catalytic activity was accompanied by altered mismatch tolerance, we evaluated Sso7d–Cas12a using guide RNAs containing defined double mismatches distributed along the spacer sequence, with particular emphasis on the PAM-proximal seed region, which is known to be critical for Cas12a target recognition and cleavage^38^ (**Fig. 6a**). Cas12a enzymes have been extensively reported to exhibit strong intolerance to mismatches within this seed region, while displaying increased tolerance toward mismatches in PAM-distal positions ^39^. Consistent with these established specificity profiles, both Sso7d–Cas12a and WT showed pronounced loss of activation when double mismatches were introduced within the seed region, while mismatches at distal positions resulted in progressively higher residual activity (**Fig. 6b**). Importantly, positions M6 and M9 showed a larger absolute increase in trans-cleavage signal for Sso7d–Cas12a than would be expected from a uniform fold-improvement over wild-type. These positions are located at the boundary between the PAM-proximal seed region and the PAM-distal region of the protospacer, where R-loop propagation is expected to be slower. The greater benefit of Sso7d at these positions is consistent with the fusion protein stabilising partially formed R-loop intermediates that stall at this boundary, rather than accelerating all productive RNP complexes equally. This also fits with the kinetic data showing that the main effect of Sso7d is on steps after initial target binding. Together with previously reported specificity characteristics of wild-type Cas12a, these data indicate that the enhanced catalytic activity of Sso7d–Cas12a is achieved without compromising sequence-dependent target discrimination.

**Figure 6.**
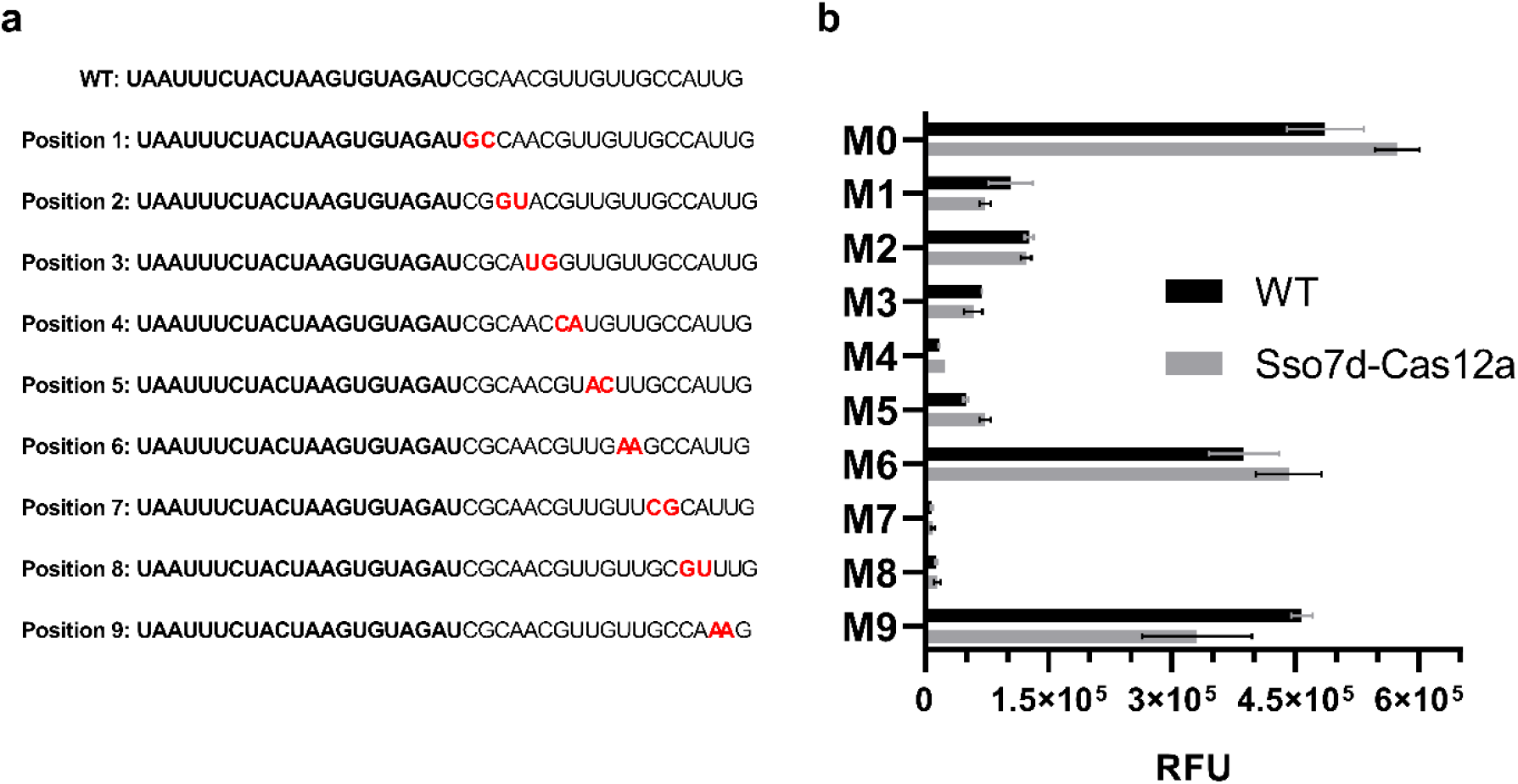
Effect of guide RNA mismatches on Cas12a activation. (a) Schematic of guide RNA 2 showing the positions of introduced double mismatches (M1-M9) along the spacer sequence, numbered relative to the PAM-proximal end of the protospacer. M0 shows the fully matched guide RNA. (b) Fluorescence signal (RFU) generated by wild-type Cas12a and Sso7d–Cas12a in the presence of target DNA programmed with the indicated double-mismatch guide RNAs. WT – Wild type; RFU – Relative fluorescent units; Reactions were performed at 37 °C for 1 h, with fluorescence recorded every 1 min. Data represent mean ± s.d. from at least three independent replicates.

We next quantified the analytical limit of detection (LOD) of the Sso7d-Cas12 fusion protein using the same standardized in vitro trans-cleavage assay. LOD measurements were performed using serial dilutions of the double-stranded target DNA and guide RNA 2, which exhibited the largest fold enhancement in activity among the guides tested. Reaction rates were extracted from the linear region of the fluorescence time courses and used as the primary metric for detection sensitivity. Across the tested concentration range, Sso7d–Cas12a exhibited a pronounced increase in reaction rate at low target DNA concentrations compared to wild-type Cas12a (**Fig. 7a**), enabling reliable discrimination from the non-template control at 0.15 nM DNA inputs (two way A-NOVA) whereas the wild type protein only produced reliable results at around 0.3 nM with on/off signal response being observed at lower concentrations. Rate-based analysis yielded a clear separation between background and target-containing reactions, while endpoint fluorescence measurements (RFU value at the 60 min timepoint) showed consistent, concentration-dependent signal accumulation (**Fig. 7b**). Importantly, the enhanced sensitivity of the fusion construct was achieved without increased background activation, indicating that improved LOD arises from accelerated productive trans-cleavage rather than elevated baseline signal. These results establish that fusion of the Sso7d DNA-binding domain significantly improves the analytical sensitivity of Cas12a in a controlled in vitro setting, providing the mechanistic foundation for subsequent application to diagnostically relevant workflows. It should be noted that, much like any other CRISPR experiment, guide RNA screening is necessary to determine which spacer offer the best targeting efficiency and that without this step the observed LOD, specificity and off target would be significantly lower.

**Figure 7.**
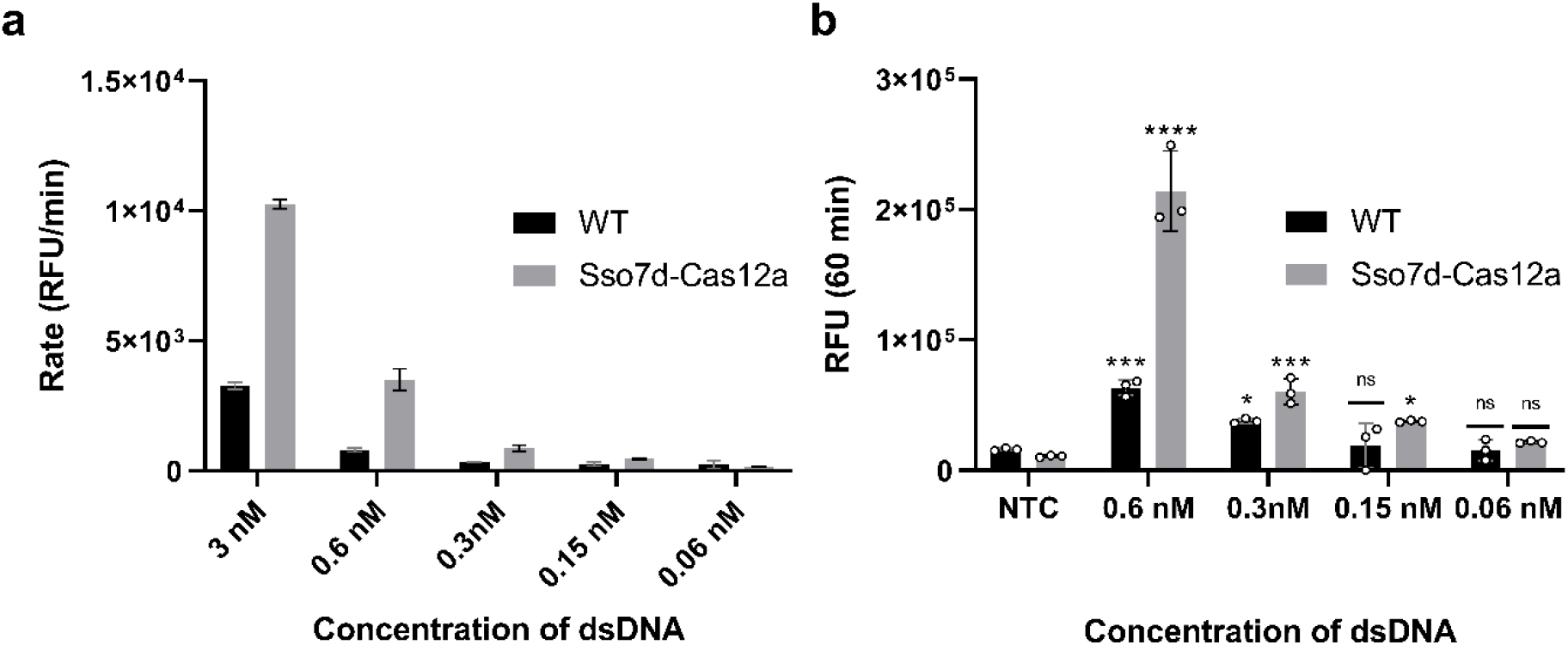
Improved limit of detection by Sso7d–Cas12a. (a) Reaction rate as a function of target DNA concentration for wild-type Cas12a and Sso7d–Cas12a. Rates were derived from the linear region of fluorescence time course. (b) Endpoint fluorescence signal (RFU) measured at 60 minutes for the corresponding target concentrations. WT – Wild type; RFU – Relative fluorescent units; NTC – non template control. Statistical significance relative to the no-template control is indicated. Reactions were performed at 37 °C for 1 h, with fluorescence recorded every 1 min. Data represent mean ± s.d. from at least three independent replicates.

### Improved analytical performance in antimicrobial resistance detection

To evaluate performance in a diagnostically relevant setting, we applied the Sso7d–Cas12a system for the detection of antimicrobial resistance (AMR) markers in de-identified clinical bacterial isolates. *Acinetobacter baumannii* (*A. baumannii*) is a Gram-negative opportunistic pathogen that is a major cause of hospital-acquired infections and is frequently associated with high levels of antimicrobial resistance (AMR). The global dissemination of multidrug-resistant *A. baumannii* has emerged as a serious public health challenge, largely driven by the horizontal acquisition of resistance determinants via mobile genetic elements, including plasmids. Following genomic DNA extraction from multiple *A. baumannii* isolates, LAMP was performed targeting the *blaOXA-51, blaOXA-23*, and *blaOXA-24* genes ^40–44^ (**Fig. 8**). LAMP reactions targeting *blaOXA-51* yielded positive amplification signals for all strains tested, consistent with the intrinsic presence of this gene in *A. baumannii* (**Fig. 8c**). In contrast, no amplification was observed for *blaOXA-23 (***Supplementary Fig. 4 and Fig. 5)** in any of the isolates under the tested conditions. Unexpectedly, LAMP reactions targeting *blaOXA-24* produced positive amplification signals across all tested strains (**Fig. 8a**), including those that did not yield detectable products by conventional PCR (**Supplementary Fig. 4 and 5**). This result indicates that LAMP enables sensitive amplification of *blaOXA-24* sequences that may be present at low abundance or exhibit sequence variation relative to PCR primer binding sites. The specificity of the reaction was assessed via melting curve analysis (**Supplementary Fig. 7**) and via sanger sequencing for both PCR and LAMP reaction (**Supplementary Table 3 and Supplementary Fig. 8**). To evaluate the ability of Sso7d-Cas12a fusion to detect *blaOXA-51* and *blaOXA-24*, the amplicons from these reactions were introduced into Cas12a trans-cleavage reactions and monitored via fluorescence output (**Fig. 8b and 8d**). Sso7d-Cas12a generated a strong and reproducible fluorescence signal in response to amplicons derived from all genomic DNA samples, demonstrating robust detection and as much 30-fold increase in the dynamic range across multiple alleles (calculated via end point background subtracted fluorescence). In contrast, reactions containing wild-type Cas12a failed to produce a significant fluorescence signal above background under identical assay conditions. Importantly, sequencing confirmed both the presence of the *blaOXA-24* and *51* in the LAMP amplified reaction (**Supplementary Table 3**) as well as that the guide RNA target sites were fully conserved across all tested isolates (**Supplementary Fig. 8**), indicating that the failure of wild-type Cas12a to generate a detectable signal was not due to target sequence mismatch. Rather, this likely reflects limitations in Cas12a activation under conditions where target DNA is structurally heterogeneous and derived from genomic amplification, as is common in clinical samples. Allelic diversity and variation in flanking sequence context can modulate R-loop formation and conformational activation without altering guide complementarity. The enhanced performance of the fusion Cas12a suggests that increased DNA engagement and stabilization of the target-bound state lowers the activation threshold required for productive trans-cleavage, enabling reliable detection across diverse clinical alleles. This clear difference highlights the enhanced functional performance of the engineered fusion protein in coupling LAMP amplification to CRISPR-based detection. These findings underscore the specificity and sensitivity of Sso7d-Cas12a within an integrated AMR detection workflow.

**Figure 8.**
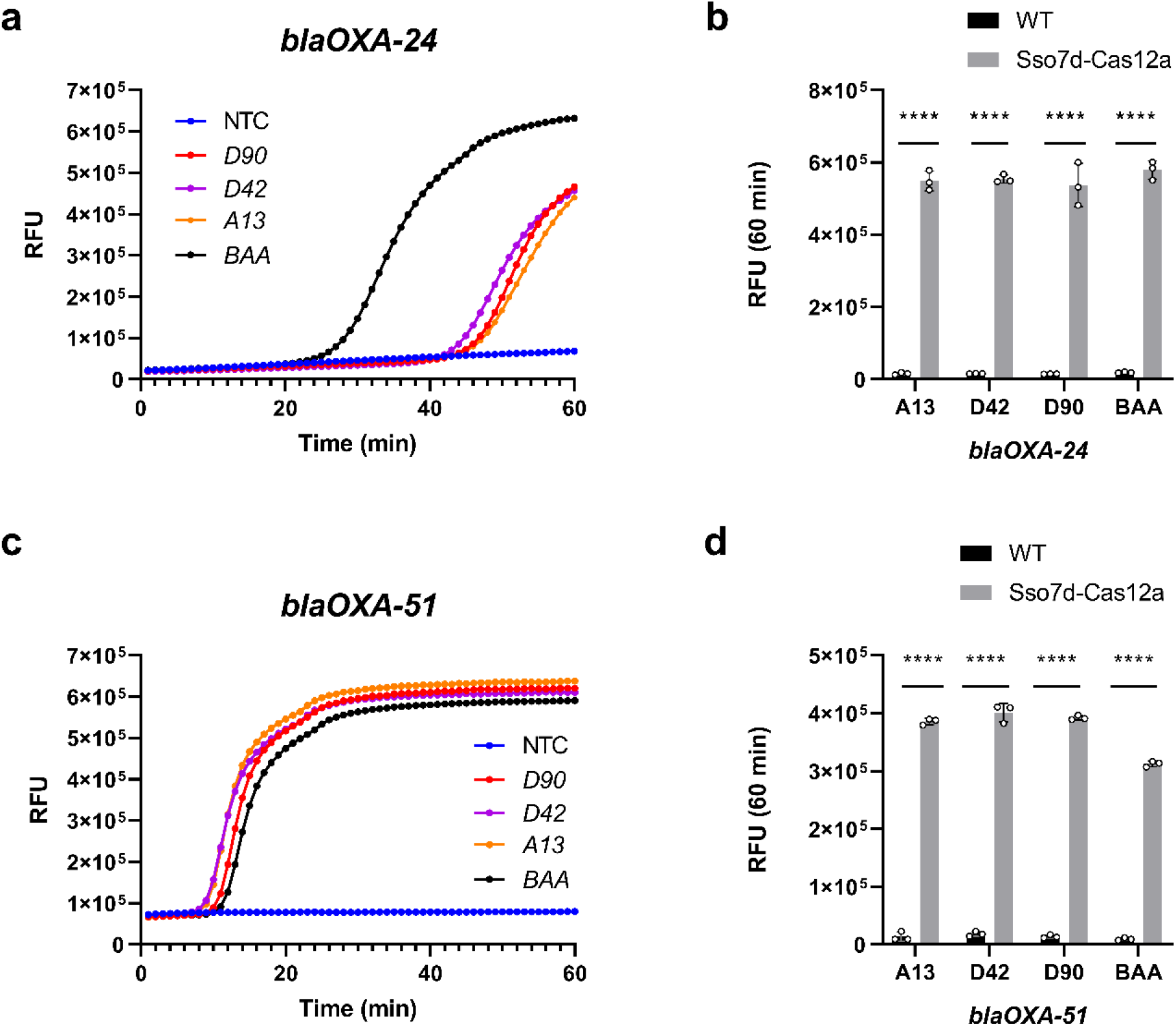
LAMP amplification and CRISPR–Cas12a detection of AMR genes from *A. baumannii* genomic DNA. (a) LAMP amplification of *blaOXA-24* from genomic DNA of multiple *A. baumannii* isolates, including strains that were negative or weakly positive by conventional PCR. (b) CRISPR–Cas12a fluorescence detection of *blaOXA-24* amplicons using Sso7d-Cas12 and wild type Cas12a. (c) LAMP amplification of *blaOXA-51*. (d) CRISPR–Cas12a fluorescence detection of *blaOXA-51* amplicons using Sso7d-Cas12 and wild type Cas12a. WT – Wild type; NTC – Non-template-control (negative); D90, D42, A13 are Japanese clinical isolates whereas BAA is BAA-1794 MDR reference strain, *A. baumannii* isolates with different (unknown) genetic profiles. Trans cleavage reactions were performed at 37 °C for 1 h, with fluorescence recorded every 1 min. LAMP reactions were performed at 65 °C for 1 h, with fluorescence recorded every 1 min. Data represent mean ± s.d. from at least three independent replicates.

## DISCUSSION

In this study, we demonstrate that fusion of the hyperthermophilic DNA-binding protein Sso7d to Cas12a substantially enhances catalytic activity, thermal robustness, and analytical performance. Across multiple assays, the Sso7d–Cas12a fusion consistently exhibited approximately 4-fold higher reaction velocities, a 4.6-fold increase in *k*_cat_, and preserved—or modestly improved—substrate affinity.

These gains were observed across multiple guide RNAs and substrate formats and were maintained across temperatures, with the magnitude of enhancement varying with guide RNA baseline activity.

The kinetic data support a model in which Sso7d accelerates the conformational activation step that converts the target-bound RNP from an inactive to a catalytically competent state. Three observations from the data are relevant. First, the main kinetic change is a 4.6-fold increase in *k*_cat_ with only a 1.46-fold increase in *K*_m_. If Sso7d acted by increasing DNA binding affinity, a decrease in *K*_m_ would be expected. The small *K*_m_ increase instead suggests a minor effect on RNP–target assembly, while the large *k*_cat_ increase points to faster cleavage once the complex is formed. Second, the improvement in trans-cleavage (4.6-fold *k*_cat_) is larger than the improvement in cis-cleavage (∼1.5-fold at 37°C). Both activities require R-loop formation, but trans-cleavage acts on the single-stranded reporter while cis-cleavage involves cutting the double-stranded target. The larger trans-cleavage gain suggests an effect on RuvC engagement with single-stranded DNA that is downstream of R-loop formation. Together, these observations are consistent with Sso7d binding to the dsDNA flanking the protospacer, stabilising the target-bound RNP, and lowering the energy barrier for the conformational switch to the active state. We speculate that the C-terminal positioning of Sso7d may occlude the PAM-interacting domain of LbCas12a or sterically hinder RuvC domain access to substrate, whereas the N-terminal positioning places Sso7d on the distal face of the protein, allowing non-specific DNA engagement without interfering with guide RNA loading or catalytic domain function.

The activity improvement does not reduce specificity. Mismatch profiling showed that Sso7d–Cas12a and wild-type Cas12a show the same pattern of sensitivity to mismatches: activity is strongly suppressed by mismatches in the seed region and more tolerant of mismatches in PAM-distal positions. This indicates that Sso7d does not change how the guide RNA reads the target sequence. The off-target data with KanR supports this: wild-type Cas12a showed significant trans-cleavage above the NTC for guide RNAs 1 and 2 in the presence of the off-target substrate, while Sso7d–Cas12a showed no significant difference from NTC under the same conditions. The improvement in off-target discrimination is consistent with Sso7d accelerating the productive on-target pathway proportionally more than any residual off-target activation.

Independent verification of antimicrobial resistance gene presence in *A. baumannii* genomic DNA provided a biologically relevant framework for evaluating the functional impact of Cas12a engineering. Although LAMP confirmed the presence of intrinsic resistance markers such as *blaOXA-51* and acquired resistance determinants including *blaOXA-24*, only Sso7d-Cas12a consistently produced a fluorescence signal above background for all isolates tested, with up to 30-fold higher endpoint signal than wild-type Cas12a. Notably, PCR failed to amplify several AMR targets despite intact primer-binding sites, highlighting the sensitivity of conventional PCR to template complexity in clinical samples. In contrast, the strand-displacing polymerase and isothermal reaction conditions employed in

LAMP conferred enhanced tolerance to DNA topology and inhibitory components, rendering it more compatible with downstream CRISPR-based detection. The failure of wild-type Cas12a to generate a detectable signal under the same conditions, despite confirmed target availability and conserved guide RNA binding sites verified by Sanger sequencing, shows that low catalytic efficiency is a limiting factor in CRISPR-based detection when template DNA is structurally complex, as is typical of LAMP amplicons.

Beyond diagnostics, the Sso7d–Cas12a architecture may also have implications for genome engineering and synthetic biology. Although genome-editing outcomes were not examined here, enhanced target engagement and accelerated activation could influence editing efficiency or the kinetics of DNA repair pathway recruitment. Conversely, the pronounced increase in trans-cleavage activity may be advantageous for high-throughput screening, molecular recording, or CRISPR-based logic and signal-processing systems. More broadly, these results suggest that fusing a nonspecific DNA-binding domain to Cas12a is a viable approach to improving catalytic performance without modifying the active site or the guide RNA. Whether alternative DNA-binding domains would produce similar effects has not been tested here, but the approach could in principle be extended to other Cas orthologs or other small DNA-binding proteins with different thermal or salt tolerances.

Despite these results, several limitations should be noted. Although the Sso7d–Cas12a fusion retained measurable activity across an extended temperature range, detection of low target DNA concentrations was not achievable at elevated temperatures. This outcome is consistent with the fact that LbaCas12a is not a thermophilic enzyme and exhibits optimal performance near physiological temperatures. Accordingly, the enhanced sensitivity observed at 37 °C likely reflects improved target engagement and catalytic efficiency within the native operating window of the Cas12a ortholog. At higher temperatures, reduced enzyme–target stability and shortened dwell times may limit productive activation events, particularly at low target abundance, despite the presence of the thermostable Sso7d domain. These results indicate that while accessory DNA-binding domain fusion can broaden the usable temperature range of Cas12a, it does not fully overcome the intrinsic thermal constraints of the underlying nuclease.

In summary, fusion of Sso7d to the N terminus of Cas12a increases *k*_cat_ 4.6-fold, shortens the time to signal plateau, and improves detection sensitivity without reducing specificity. The kinetic data point to an effect on the conformational activation step after target binding rather than on initial DNA recognition. This approach is straightforward to implement and may be applicable to other Cas orthologs.

## MATERIAL AND METHODS

### DNA CLONING

The LbCpf1-2NLS plasmid was a gift from Jennifer Doudna (Addgene plasmid #102566; http://n2t.net/addgene:102566 ; RRID:Addgene_102566). The Sso7d *E. coli*. codon optimized sequence was synthesized by ThermoFisher as dsDNA. DNA oligonucleotides were purchased from Hokkaido System Science and/or ThermoFisher. KOD Plus Neo (Toyobo Life Science) was used for the PCR amplification. PCR products were purified by agarose gel electrophoresis using the FastGene™ Gel / PCR Extraction Kit (Nippon Genetics, FG-91302). Small-scale plasmid DNA was obtained from a 1.5 mL LB-liquid bacterial culture by alkaline lysis and ethanol precipitation. DNA sequencing of the cDNA constructs was performed using the BigDye Terminator v1.1 Cycle Sequencing kit (Life Technologies). All DNA and amino acid sequences are available in the supplementary information file.

### GUIDE RNA SYNTHESIS AND PURIFICATION

gRNAs (**Supplementary Table 1**) were synthesized using the HiScribe ® T7 High Yield RNA Synthesis Kit. First, we made a fill-in PCR to generate dsDNA with a T7 promoter for transcription. After this, the dsDNA was purified using a column purification kit (Nippon Genetics, FG-91302). Next, the HiScribe ® T7 High Yield RNA Synthesis Kit components were assembled as recommended by the manufacturer, incubated at 37 ºC with shaking o.n. For purification of the gRNA, 30 min Dnase I treatment at 37 ºC was required to remove dsDNA template. Next, phenol chloroform extraction was used, followed by ethanol precipitation. After drying at 55 ºC, the pellet was resuspended in 50 µL 0.1mM EDTA and quantified using nanodrop and 2% agarose gel.

### PROTEIN PURIFICATION

E. coli strain Rosetta 2 (DE3) was transformed with the pBR322 plasmid harbouring Cas12a variants fused with an N-terminal Polyhistidine tag (**Supplementary Table 2**) by the heat shock method at 42 °C for 45 s. Transformants were spread on an LB plate containing 100 µg/mL carbenicillin and 33 µg/mL chloramphenicol and incubated at 37 °C overnight. Three, 2 ml pre-cultures were prepared in terrific broth (TB) with carbenicillin/chloramphenicol. These were incubated overnight at 37 °C. After incubation, a 200 mL TB culture was inoculated with 1 mL of the pre-culture mixture and grown at 37 °C until OD 600∼0.8 (usually around 4 to 5h). Before induction, cultures were incubated on ice for 30 min. After this, 80 µL of 1M IPTG was added, and flasks were incubated overnight at 18 ºC. Cells were collected by centrifugation at 8,000 rpm for 20 min, and the pellet was resuspended in 10 mL lysis buffer with protease inhibitors and benzonase nuclease. Cells were disrupted via sonication and then centrifuged at 14,000 rpm for 30 min. The supernatant was then filtered using a 0.22 µm filter and added to the Ni-NTA agarose affinity columns (Qiagen). After this, the resin was washed with 20 mM imidazole (FUJIFILM Wako) and eluted with 230 mM imidazole in 500 µL fractions after washing the column multiple times with high ionic strength buffer (2 M NaCl). Finally, the buffer was exchanged to 20 mM HEPES (pH 7.4), 300 mM KCl, 1 mM DTT, 10% Glycerol using PD-10 column (Cytiva). The resulting protein was then further purified using an Chromaster System (Hitachi-Hightech) equipped with a Superdex 200 Increase 10/300 GL (Cytiva). Sso7d-Cas12 solution (50 μL) was loaded onto the column, filtered through a 0.20 μm pore filter (Millipore). A 20 mM HEPES buffer containing 300 mM NaCl was used for the mobile phase at a flow rate of 0.50 mL/min. Absorption was recorded at 280 nm (**Supplementary Fig. 1**). After purification, sodium dodecyl sulfate–polyacrylamide gel electrophoresis (SDS-PAGE) was performed to determine the degree of purification (**Supplementary Fig. 2**). Pooled protein fractions were concentrated using an Amicon filter MWCO 30K. All protein fractions were stored in 20 mM HEPES (pH 7.4), 300 mM KCl, 1 mM DTT, 10% Glycerol until further use. Amino acid sequence is detailed in **Supplementary Table 2**.

### CRISPR ASSAY

For the CRISPR-based detection system, we used commercially available EnGenLbaCas12a (NEB) and in-house purified Sso7d-Cas12. Firstly, Cas12-gRNA complexes were generated by incubating LbaCas12a (200 nM, final concentration) with the specific gRNA (400 nM final concentration for 30 min at 37ºC in or 10 mM Tris-HCl, 100 mM NaCl, 10 mM MgCl_2_, 100 µg/ml BSA pH 9 (for Sso7d-Cas12a) in a final volume of 25 µL. After incubation, complexes were diluted to 30 nM and the ssDNA fluorescent reporter molecule was added at a final concentration of 1 µM and placed on ice. 9 µL of complexes were added to 1 µL of target DNA (**Supplementary Table 3**) or water (NTC) in a 48-well plate. Fluorescence signal was measured every minute for 1 h using the FAM channel (495ex/520em) in an Applied Biosystems StepOne Real-Time PCR system.

### IN SILICO ANALYSIS

Target DNA sequences were aligned using Clustal Omega using available genomes from GenBank (NCBI). The LAMP primers (**Supplementary Table 2)** were designed (*blaOXA-24*) via PrimerExplorer v.5 (https://primerexplorer.jp/e/) or based on previously published primers (*blaOXA51*^45^, *blaOXA-23*^46^). Guide RNAs were manually designed and screened for proof-of-concept tests (AmpR detection, gRNA 1, 2, 3, 4) and AMR (*blaOXA-51, blaOXA-24*) or based on previously published sequences (*blaOXA-23*^47^).

### KINETIC ANALYSIS

Trans-cleavage activity was measured using a fluorophore–quencher–labeled single-stranded DNA reporter. For steady-state reporter kinetics, Cas12a RNPs were activated by incubation with target DNA (from 300 nM to 0.06 nM). Fluorescence was monitored in real time using a plate reader with appropriate excitation and emission settings (SYBR green channel). Initial reaction velocities were calculated from the linear region of the fluorescence traces (R^2^=0.99) when the fluorescence signal increase was in the linear range, by linear regression with a linear function of:

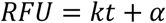

where α is a zero-time fluorescence intensity, k is the slope (RFU/min) representing the initial rate for trans nuclease activity when the reaction was in the steady state and t is time. These ratios were converted to molar cleavage rates using a calibration curve generated with a fully cleaved reporter. Michaelis–Menten parameters (*k*_cat_ and *K*_m_) were obtained by nonlinear regression of initial velocity versus DNA concentration.

t_sat_ was defined as the time at which the fluorescence signal reached plateau. Fluorescence traces were fitted to a one-phase association equation:

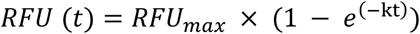

Using nonlinear regression in GraphPad Prism. t_sat_ was defined as the time at which the fitted curve reached 95% of the asymptotic maximum (RFU_max_).

### LAMP ASSAYS

10× LAMP primer mixtures were assembled using a final concentration of 0.2 µM of F3/B3, 0.4 µM of LF/LB, and 1.6 µM of FIP/BIP in nuclease-free water, this was added to the 2× WarmStart Colorimetric LAMP 2× Master Mix. Melt curve analysis was performed from 35 to 95 °C with a heating ramp of 0.3 °C/s. LAMP primers have been previously validated as highly specific with minimal cross reactivity as confirmed by the in-silico analysis.

### ANALYTICAL VALIDATION (LOD)

LoD is the lowest detectable concentration at which around 95% of all true positive replicates test positive. The analytical validation was performed by using duplex DNA that was amplified and purified in-house. From this stock, serial dilutions were made until the detection limit was reached (lowest concentration at which 3 of the replicates produce signal significantly above the non-template control reaction).

### CLINICAL VALIDATION (AMR)

The Acinetobacter baumannii clinical isolates used in this study were fully de-identified legacy strains obtained from a previously published study ^48^. Under the national Ethical Guidelines for Medical and Biological Research Involving Human Subjects (Ministry of Health, Labour and Welfare / Ministry of Education, Culture, Sports, Science and Technology, Japan), ethics approval and informed consent were deemed unnecessary by the Research Ethics Committee of SANKEN (The Institute of Scientific and Industrial Research), The University of Osaka, Japan as the isolates constitute pre-existing, irreversibly de-identified specimens collected for a prior study and used here for secondary research purposes. All methods were carried out in accordance with relevant guidelines and regulations. Genomic DNA was extracted using a previously described boiling method. After overnight culture in LB at 37 ºC, cells were pelleted, washed with PBS buffer, followed by the resuspension of the pellet in TE buffer (100 µL). Samples were then heated at 100 ºC for 5 min, followed by centrifugation. The supernatant was either used directly for DNA detection via LAMP or further purified via chloroform extraction for PCR.

### STATISTICAL INFORMATION

Data were analysed using GraphPad Prism version 8.0.0 for Windows, GraphPad Software, San 473 Diego, California USA, www.graphpad.com. Statistical significance was determined by two-way A-NOVA, and all data were shown as mean ± s.d. Asterisks indicate * p ≤ 0.05; ** p<0.01; *** p<0.001; **** p<0.0001, and “ns” is non-significant.

## DATA AVAILABILITY

The datasets used and/or analysed during the current study are available from the corresponding author on reasonable request.

## ACKNOWLEDGEMENTS

We thank Dr. Shigemori for his advice on the characterization of fusion Cas12a proteins.

## SUPPLEMENTARY DATA

Supplementary Data are available online.

