## Supplementary for "Fusion of a non-specific DNA-binding domain enhances Cas12a trans-cleavage for robust nucleic-acid diagnostics"

**SUPPLEMENTARY INFORMATION**

This Supplementary Information file accompanies the manuscript entitled: “Fusion of a non-specific DNA-binding domain enhances Cas12a trans-cleavage for robust nucleic-acid diagnostics"

**Supplementary Table 1 |** List of all crRNAs used in this study, including target name, sequence, and protospacer adjacent motif (PAM).

| gRNA ID | Target gene | Sequence (5’-3’) | PAM |
|---|---|---|---|
| gRNA-1 | AmpR | **UAAUUUCUACUAAGUGUAGAU**GUAUGGCUUCAUUCAGCUCCG | **TTTG** |
| gRNA-2 | AmpR | **UAAUUUCUACUAAGUGUAGAU**CGCAACGUUGUUGCCAUUG | **TTTG** |
| gRNA-3 | AmpR | **UAAUUUCUACUAAGUGUAGAU**UCCGCCUCCAUCCAGUCUAU | **TTTA** |
| gRNA-4 | AmpR | **UAAUUUCUACUAAGUGUAGAU**UUGCUGAUAAAUCUGGAGCCGG | **TTTA** |
| gRNA-OXA23 | blaOXA-23 | **UAAUUUCUACUAAGUGUAGAU**CAUGAGAUCAAGACCGAUACG | **TTTG** |
| gRNA-OXA24 | blaOXA-24 | **UAAUUUCUACUAAGUGUAGAU**GGUGAGGCAAUGGCAUUGUC | **TTTA** |
| gRNA-OXA51 | blaOXA-51 | **UAAUUUCUACUAAGUGUAGAU**GAAGGUCGAAGCAGGUACA | **TTTT** |

**Supplementary Table 2 |** Amino acid sequence of Sso7d-Cas12a

| >Sso7d-Cas12 |
|---|
| MATVKFKYKGEEKEVDISKIKKVWRVGKMISFTYDEGGGKTGRGAVSEKDAPKELLQMLEKQKKSGSE<br>TPGTSESATPESSKLEKFTNCYSLSKTLRFKAIPVGKTQENIDNKRLLVEDEKRAEDYKGVKKLLDRYYL<br>SFINDVLHSIKLKNLNNYISLFRKKTRTEKENKELENLEINLRKEIAKAFKGNEGYKSLFKKDIIETILPEFL<br>DDKDEIALVNSFNGFTTAFTGFFDNRENMFSEEAKSTSIAFRCINENLTRYISNMDIFEKVDAIFDKHEVQ<br>EIKEKILNSDYDVEDFFEGEFFNFVLTQEGIDVYNAIIGGFVTESGEKIKGLNEYINLYNQKTKQKLPKFKP<br>LYKQVLSDRESLSFYGEGYTSDEEVLEVFRNTLNKNSEIFSSIKKLEKLFKNFDEYSSAGIFVKNGPAISTIS<br>KDIFGEWNVIRDKWNAEYDDIHLKKKAVVTEKYEDDRRKSFKKIGSFSLEQLQEYADADLSVVEKLKEI<br>IIQKVDEIYKVYGSSEKLFDADFVLEKSLKKNDAVVAIMKDLLDSVKSFENYIKAFFGEGKETNRDESFY<br>GDFVLAYDILLKVDHIYDAIRNYVTQKPYSKDKFKLYFQNPQFMGGWDKDKETDYRATILRYGSKYYL<br>AIMDKKYAKCLQKIDKDDVNGNYEKINYKLLPGPNKMLPKVFFSKKWMAYYNPSEDIQKIYKNGTFKK<br>GDMFNLNDCHKLIDFFKDSISRYPKWSNAYDFNFSETEKYKDIAGFYREVEEQGYKVSFESASKKEVDK<br>LVEEGKLYMFQIYNKDFSDKSHGTPNLHTMYFKLLFDENNHGQIRLSGGAELFMRRASLKKEELVVHPA<br>NSPIANKNPDNPKKTTTLSYDVYKDKRFSEDQYELHIPIAINKCPKNIFKINTEVRVLLKHDDNPYVIGIDR<br>GERNLLYIVVVDGKGNIVEQYSLNEIINNFNGIRIKTDYHSLLDKKEKERFEARQNWTSIENIKELKAGYI<br>SQVVHKICELVEKYDAVIALEDLNSGFKNSRVKVEKQVYQKFEKMLIDKLNYMVDKKSNPCATGGALK<br>GYQITNKFESFKSMSTQNGFIFYIPAWLTSKIDPSTGFVNLLKTKYTSIADSKKFISSFDRIMYVPEEDLFEF<br>ALDYKNFSRTDADYIKKWKLYSYGNRIRIFRNPKKNNVFDWEEVCLTSAYKELFNKYGINYQQGDIRAL<br>LCEQSDKAFYSSFMALMSLMLQMRNSITGRTDVDFLISPVKNSDGIFYDSRNYEAQENAILPKNADANG<br>AYNIARKVLWAIGQFKKAEDEKLDKVKIAISNKEWLEYAQTSVKH |

**Supplementary Table 3 |** DNA substrates used for cis-cleavage, trans-cleavage activation, and LOD measurements.

| Description | Sequence (5'-3') |
|---|---|
| ssDNA reporter probe | 5' FITC (6-FAM) - TTATTATTATT - 3' BHQ1 |
| Linear dsDNA (AmpR) | ATGAGTATTCAACATTTCCGTGTCGCCCTTATTCCCTTTTTTGCGGCATTTTGCCTTC<br>CTGTTTTTGCTCACCCAGAAACGCTGGTGAAAGTAAAAGATGCTGAAGATCAGTTG<br>GGTGCACGAGTGGGTTACATCGAACTGGATCTCAACAGCGGTAAGATCCTTGAGA<br>GTTTTCGCCCCGAAGAACGTTTTCCAATGATGAGCACTTTTAAAGTTCTGCTATGTG<br>GCGCGGTATTATCCCGTATTGACGCCGGGCAAGAGCAACTCGGTCGCCGCATACA<br>CTATTCTCAGAATGACTTGGTTGAGTACTCACCAGTCACAGAAAAGCATCTTACGG<br>ATGGCATGACAGTAAGAGAATTATGCAGTGCTGCCATAACCATGAGTGATAACAC<br>TGCGGCCAACTTACTTCTGACAACGATCGGAGGACCGAAGGAGCTAACCGCTTTTT<br>TGCACAACATGGGGGATCATGTAACTCGCCTTGATCGTTGGGAACCGGAGCTGAA<br>TGAAGCCATACCAAACGACGAGCGTGACACCACGATGCCTGTAGCAATGGCAACA<br>ACGTTGCGCAAACTATTAACTGGCGAACTACTTACTCTAGCTTCCCGGCAACAATT<br>AATAGACTGGATGGAGGCGGATAAAGTTGCAGGACCACTTCTGCGCTCGGCCCTT<br>CCGGCTGGCTGGTTTATTGCTGATAAATCTGGAGCCGGTGAGCGTGGGTCTCGCGG<br>TATCATTGCAGCACTGGGGCCAGATGGTAAGCCCTCCCGTATCGTAGTTATCTACA<br>CGACGGGGAGTCAGGCAACTATGGATGAACGAAATAGACAGATCGCTGAGATAG<br>GTGCCTCACTGATTAAGCATTGGTAA |
| Circular dsDNA | GATCTCGATCCCGCGAAATTAATACGACTCACTATAGGGAGACCACAACGGTTTCC<br>CTCTAGAAATAATTTTGTTTAACTTTAAGAAGGAGATATACATATGCGGGGTTCTC<br>ATCATCATCATCATCATGGTATGGCTAGCATGACTGGTGGACAGCAAATGGGTCGG<br>GATCTGTACGACGATGACGATAAGGATCCGAGCTCGAGATCTGCAGCTGGTACCA<br>TGGAATTCGAAGCTTGATCCGGCTGCTAACAAAGCCCGAAAGGAAGCTGAGTTGG |

| | |
|---|---|
| | CTGCTGCCACCGCTGAGCAATAACTAGCATAACCCCTTGGGGCCTCTAAACGGGTC<br>TTGAGGGGTTTTTTGCTGAAAGGAGGAACTATATCCGGATCTGGCGTAATAGCGAA<br>GAGGCCCGCACCGATCGCCCTTCCCAACAGTTGCGCAGCCTGAATGGCGAATGGG<br>ACGCGCCCTGTAGCGGCGCATTAAGCGCGGCGGGTGTGGTGGTTACGCGCAGCGT<br>GACCGCTACACTTGCCAGCGCCCTAGCGCCCGCTCCTTTCGCTTTCTTCCCTTCCTT<br>TCTCGCCACGTTCGCCGGCTTTCCCCGTCAAGCTCTAAATCGGGGGCTCCCTTTAG<br>GGTTCCGATTTAGTGCTTTACGGCACCTCGACCCCAAAAAACTTGATTAGGGTGAT<br>GGTTCACGTAGTGGGCCATCGCCCTGATAGACGGTTTTTCGCCCTTTGACGTTGGA<br>GTCCACGTTCTTTAATAGTGGACTCTTGTTCCAAACTGGAACAACACTCAACCCTA<br>TCTCGGTCTATTCTTTTGATTTATAAGGGATTTTGCCGATTTCGGCCTATTGGTTAA<br>AAAATGAGCTGATTTAACAAAAATTTAACGCGAATTTTAACAAAATATTAACGCTT<br>ACAATTTAGGTGGCACTTTTCGGGGAAATGTGCGCGGAACCCCTATTTGTTTATTTT<br>TCTAAATACATTCAAATATGTATCCGCTCATGAGACAATAACCCTGATAAATGCTT<br>CAATAATATTGAAAAAGGAAGAGTATGAGTATTCAACATTTCCGTGTCGCCCTTAT<br>TCCCTTTTTTGCGGCATTTTGCCTTCCTGTTTTTGCTCACCCAGAAACGCTGGTGAA<br>AGTAAAAGATGCTGAAGATCAGTTGGGTGCACGAGTGGGTTACATCGAACTGGAT<br>CTCAACAGCGGTAAGATCCTTGAGAGTTTTCGCCCCGAAGAACGTTTTCCAATGAT<br>GAGCACTTTTAAAGTTCTGCTATGTGGCGCGGTATTATCCCGTATTGACGCCGGGC<br>AAGAGCAACTCGGTCGCCGCATACACTATTCTCAGAATGACTTGGTTGAGTACTCA<br>CCAGTCACAGAAAAGCATCTTACGGATGGCATGACAGTAAGAGAATTATGCAGTG<br>CTGCCATAACCATGAGTGATAACACTGCGGCCAACTTACTTCTGACAACGATCGGA<br>GGACCGAAGGAGCTAACCGCTTTTTTGCACAACATGGGGGATCATGTAACTCGCCT<br>TGATCGTTGGGAACCGGAGCTGAATGAAGCCATACCAAACGACGAGCGTGACACC<br>ACGATGCCTGTAGCAATGGCAACAACGTTGCGCAAACTATTAACTGGCGAACTAC<br>TTACTCTAGCTTCCCGGCAACAATTAATAGACTGGATGGAGGCGGATAAAGTTGCA<br>GGACCACTTCTGCGCTCGGCCCTTCCGGCTGGCTGGTTTATTGCTGATAAATCTGG<br>AGCCGGTGAGCGTGGGTCTCGCGGTATCATTGCAGCACTGGGGCCAGATGGTAAG<br>CCCTCCCGTATCGTAGTTATCTACACGACGGGGAGTCAGGCAACTATGGATGAACG<br>AAATAGACAGATCGCTGAGATAGGTGCCTCACTGATTAAGCATTGGTAACTGTCA<br>GACCAAGTTTACTCATATATACTTTAGATTGATTTAAAACTTCATTTTTAATTTAAA<br>AGGATCTAGGTGAAGATCCTTTTTGATAATCTCATGACCAAAATCCCTTAACGTGA<br>GTTTTCGTTCCACTGAGCGTCAGACCCCGTAGAAAAGATCAAAGGATCTTCTTGAG<br>ATCCTTTTTTTCTGCGCGTAATCTGCTGCTTGCAAACAAAAAAACCACCGCTACCA<br>GCGGTGGTTTGTTTGCCGGATCAAGAGCTACCAACTCTTTTTCCGAAGGTAACTGG<br>CTTCAGCAGAGCGCAGATACCAAATACTGTTCTTCTAGTGTAGCCGTAGTTAGGCC<br>ACCACTTCAAGAACTCTGTAGCACCGCCTACATACCTCGCTCTGCTAATCCTGTTA<br>CCAGTGGCTGCTGCCAGTGGCGATAAGTCGTGTCTTACCGGGTTGGACTCAAGACG<br>ATAGTTACCGGATAAGGCGCAGCGGTCGGGCTGAACGGGGGGTTCGTGCACACAG<br>CCCAGCTTGGAGCGAACGACCTACACCGAACTGAGATACCTACAGCGTGAGCTAT<br>GAGAAAGCGCCACGCTTCCCGAAGGGAGAAAGGCGGACAGGTATCCGGTAAGCG<br>GCAGGGTCGGAACAGGAGAGCGCACGAGGGAGCTTCCAGGGGGAAACGCCTGGT<br>ATCTTTATAGTCCTGTCGGGTTTCGCCACCTCTGACTTGAGCGTCGATTTTTGTGAT<br>GCTCGTCAGGGGGGCGGAGCCTATGGAAAAACGCCAGCAACGCGGCCTTTTTACG<br>GTTCCTGGCCTTTTGCTGGCCTTTTGCTCACATGTTCTTTCCTGCGTTATCCCCTGAT<br>TCTGTGGATAACCGTATTACCGCCTTTGAGTGAGCTGATACCGCTCGCCGCAGCCG<br>AACGACCGAGCGCAGCGAGTCAGTGAGCGAGGAAGCGGAAGAGCGCCCAATACG<br>CAAACCGCCTCTCCCCGCGCGTTGGCCGATTCATTAATGCAG |
| ssDNA | ATGCCTGTAGCAATGGCAACAACGTTGCGCAAACTATTAACTGG |
| Off target (NeoR/KanR) | ATGATTGAACAAGATGGATTGCACGCAGGTTCTCCGGCCGCTTGGGTGGAGAGGC<br>TATTCGGCTATGACTGGGCACAACAGACAATCGGCTGCTCTGATGCCGCCGTGTTC<br>CGGCTGTCAGCGCAGGGGCGCCCGGTTCTTTTTGTCAAGACCGACCTGTCCGGTGC<br>CCTGAATGAACTGCAAGACGAGGCAGCGCGGCTATCGTGGCTGGCCACGACGGGC<br>GTTCCTTGCGCAGCTGTGCTCGACGTTGTCACTGAAGCGGGAAGGGACTGGCTGCT<br>ATTGGGCGAAGTGCCGGGGCAGGATCTCCTGTCATCTCACCTTGCTCCTGCCGAGA<br>AAGTATCCATCATGGCTGATGCAATGCGGCGGCTGCATACGCTTGATCCGGCTACC |

| | |
|---|---|
| | TGCCCATTCGACCACCAAGCGAAACATCGCATCGAGCGAGCACGTACTCGGATGG<br>AAGCCGGTCTTGTCGATCAGGATGATCTGGACGAAGAGCATCAGGGGCTCGCGCC<br>AGCCGAACTGTTCGCCAGGCTCAAGGCGAGCATGCCCGACGGCGAGGATCTCGTC<br>GTGACCCATGGCGATGCCTGCTTGCCGAATATCATGGTGGAAAATGGCCGCTTTTC<br>TGGATTCATCGACTGTGGCCGGCTGGGTGTGGCGGACCGCTATCAGGACATAGCGT<br>TGGCTACCCGTGATATTGCTGAAGAGCTTGGCGGCGAATGGGCTGACCGCTTCCTC<br>GTGCTTTACGGTATCGCCGCTCCCGATTCGCAGCGCATCGCCTTCTATCGCCTTCTT<br>GACGAGTTCTTCTGA |
| *blaOXA-51* (sanger sequencing) | GCCAAACTCAACAAAGCTATGGTAATGATCTTGCTCGTGCTTCGACCGAGTATGTA<br>CCTGCTTCGACCTTCAAAATGCTTAATGCTTTGATCGGCCTTGAGCACCATAAGGC<br>AACCACCACAGAAGTATTTAAGTGGGATGGTAAAAAAAGGTTATTCCCAGAATGG<br>GAAAAGGACATGACCCTAGGCGATGCCATGAAAGCTTCCGCTATTCCG |
| *blaOXA-24* (sanger sequencing) | GATTTTCAAATGGGATGGTAAAAAAAGAACTTATCCTATGTGGGAGAAAGATATG<br>ACTTTAGGTGAGGCAATGGCATTGTCAGCAGTTCCAGTATATCAAGAGCTTGCAAG<br>ACGGACTGGCCTAGAGCTAATGCAGAAAGAAGTAAAGCGGGTTAATTTTGGAAAT<br>ACAAATATTGGAACACAGGTCGATA |

**Supplementary Table 4 |** Primers and oligos used in this study

| Name | Description | Sequence (5'-3') |
|---|---|---|
| F-AmpR | PCR | CGGAACCCCTATTTGTTTATTTTTC |
| R-AmpR | PCR | AAGGGATTTTGGTCATGAGATTATC |
| F-KanR | PCR | GACTAATTTTTTTATTTATGCAGAGGC |
| R-KanR | PCR | CCAACCTTTCATAGAAGGCG |
| F-OXA23 | PCR | GGGCGAGAAAAGGTCATT |
| R-OXA23 | PCR | ACCAACCAGAAATTATCAACC |
| F-OXA24 | PCR | AGATTTTCAAATGGGATGGTAA |
| R-OXA24 | PCR | ATCGACCTGTGTTCCAAT |
| F-OXA51 | PCR | GCCAAACTCAACAAAGCTATGG |
| R-OXA51 | PCR | CGGAATAGCGGAAGCTTTCA |
| F3-OXA23 | LAMP | GGGCGAGAAAAGGTCATT |
| B3-OXA23 | LAMP | ACCAACCAGAAATTATCAACC |
| FIP-OXA23 | LAMP | TAGACTGGGACTGCAGAAAGCCCGCTTGGGAAAAAGACA |
| BIP-OXA23 | LAMP | CAGGAACTTGCGCGACGTATCAATTTCAGCATTACCGAAAC |
| LB-OXA23 | LAMP | GGTCTTGATCTCATGCAAAAAGAAG |
| LF-OXA23 | LAMP | TCATGGCTTCTCCTAGTGTCA |
| F3-OXA24 | LAMP | AGATTTTCAAATGGGATGGTAA |
| B3-OXA24 | LAMP | ATCGACCTGTGTTCCAAT |
| FIP-OXA24 | LAMP | AGCAGTTCCAGTATATCAAGAGCTTTTCCAAAATTAACCCGCTT |
| BIP-OXA24 | LAMP | GACAATGCCATTGCCTCACCAAAAAGAACTTATCCTATGTGGGA |
| F3-OXA51 | LAMP | CTTATATAGTGACTGCTAATCCAA |
| B3-OXA51 | LAMP | ATTAAGCATTTTGAAGGTCGA |
| FIP-OXA51 | LAMP | ACCCGTAGTGTGTACTTCGTTAAATTTTTACAGCGCTTCAAAATCTGA |
| BIP-OXA51 | LAMP | TTAGTTATCCAACAAGGCCAAACTTTTTAGCAGGTACATACTCGGTC |
| LB-OXA51 | LAMP | AAAGCTATGGTAATGATCTTGCTCG |
| ssDNA probe | Trans cleavage assays | 5' FITC (6-FAM) - TTATTATTATT - 3' BHQ1 |

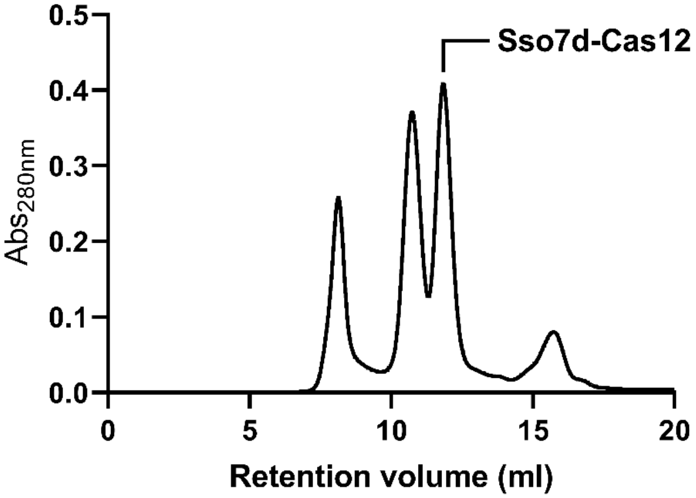


**Supplementary Figure 1 |** Size-exclusion chromatography analysis of purified Cas12a proteins to assess the purity and homogeneity. After affinity chromatography (Ni-NTA), gel filtration was used to obtain a >90% protein fraction. Since Sso7d binds DNA non-specifically, benzonase, as well as high salt washes, were used to remove contaminating nucleic acids.

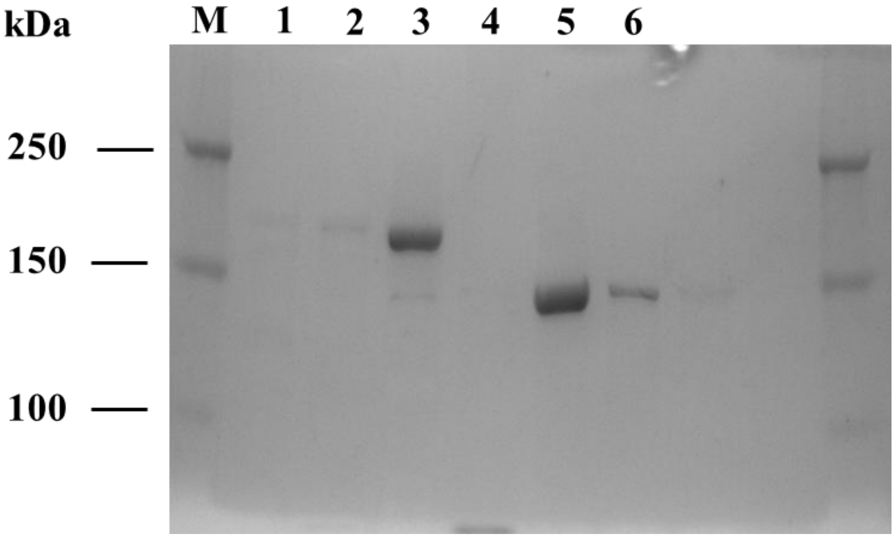


**Supplementary Figure 2 |** Uncropped 5% SDS-PAGE gel showing the purification of N Sso7d-Cas12 (Lane 5 and 6) as well as the MBP tagged wt protein (Lanes 1-4). After affinity chromatography (Ni-NTA), gel filtration was used to obtain a >90% protein fraction. Since Sso7d binds DNA non-specifically, benzonase, as well as high salt washes were used to remove contaminating nucleic acids.

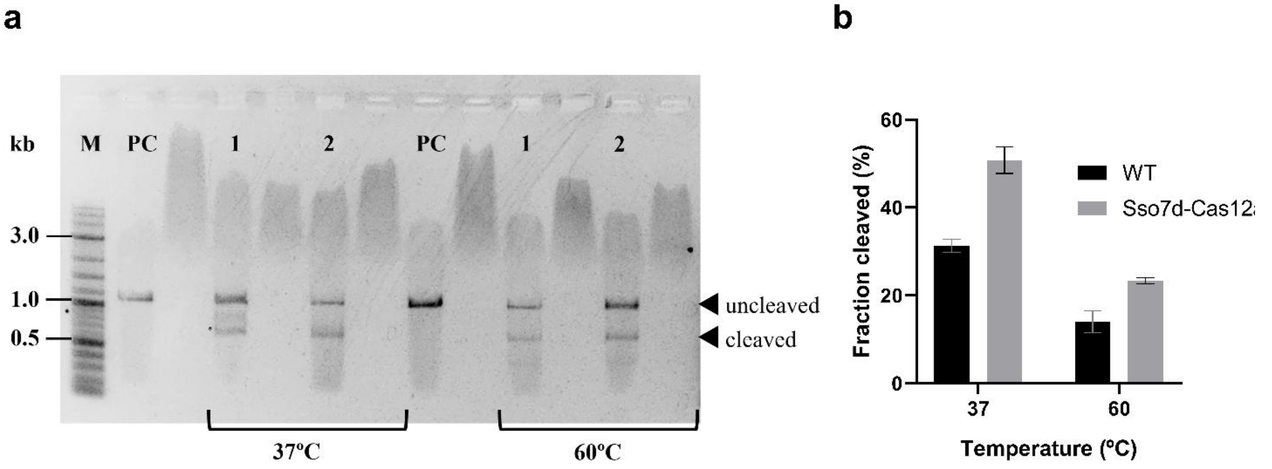


**Supplementary Figure 3 |** Quantification of cis cleavage activity. (a) Uncropped 1% Agarose gels used for cis-cleavage quantification. Quantification was made using ImageJ (Fiji) by analysing the intensities of the uncleaved vs cleaved DNA bands. (b) Cleaved DNA fraction at 37ºC and 60ºC. M: DNA Marker; PC: positive control (DNA only). Lane 1 corresponds to WT protein whereas Lane 2 corresponds to Sso7d-Cas12a.

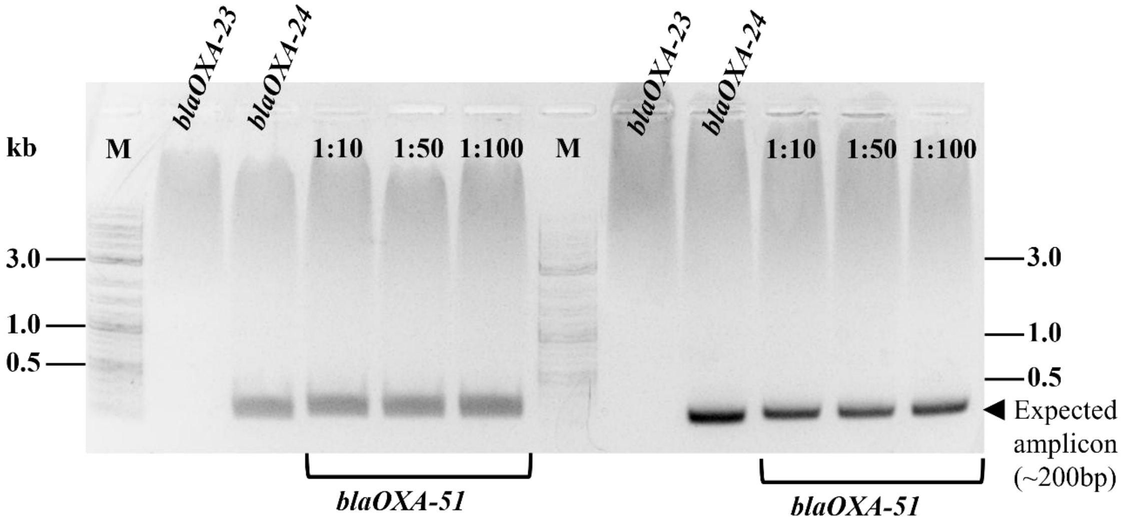


**Supplementary Figure 4 |** Uncropped agarose gel (1%) showing the optimization of PCR amplification conditions of AMR genes for reference strain BAA-1794. 2 different polymerases were used (KOD-Plus-One and Takara Ex-Taq HS) as well as 3 different DNA concentrations (400 ng – 1:10, 80 ng – 1:50 and 40 ng – 1:100 of extracted high molecular weight DNA).

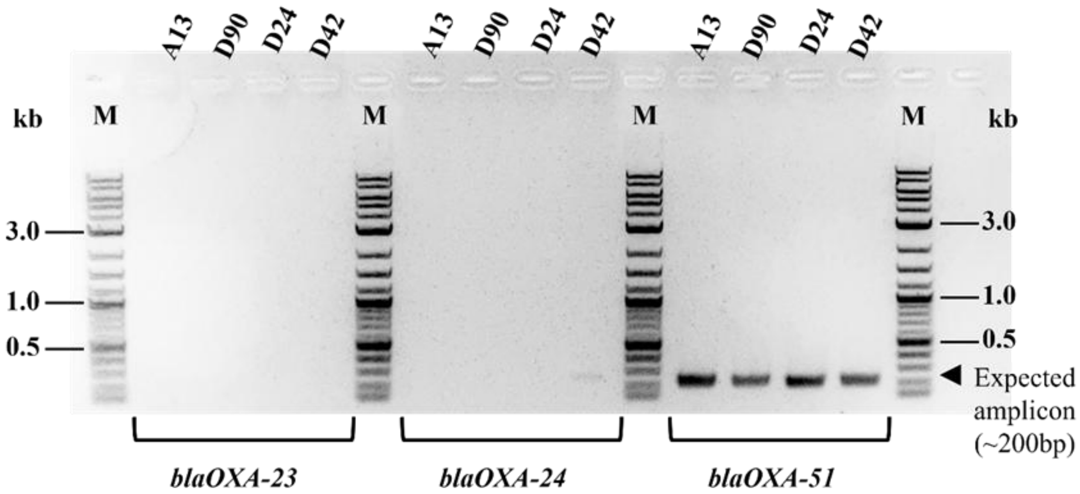


**Supplementary Figure 5 |** Uncropped agarose gel (1%) showing the PCR amplification of AMR genes. Optimal conditions were chosen, 400ng of extracted high molecular weight DNA was used to amplify the different AMR genes.

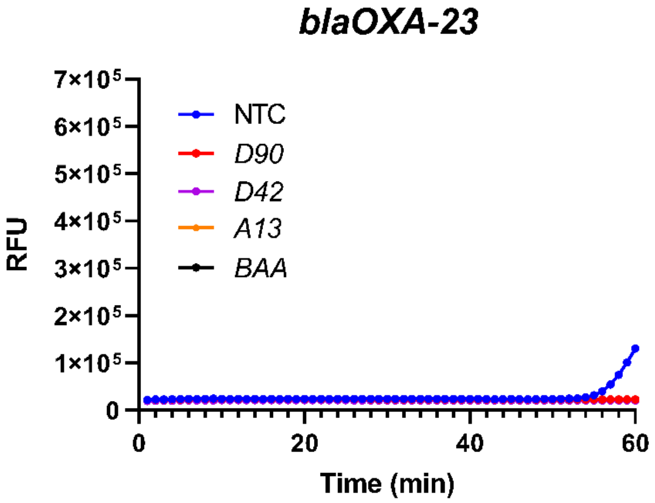


**Supplementary Figure 6 |** LAMP amplification and CRISPR–Cas12a detection of *blaOXA-23* AMR gene from A. baumannii genomic DNA. LAMP amplification was not verified for any of the strains that were tested, the results were in agreement with PCR.

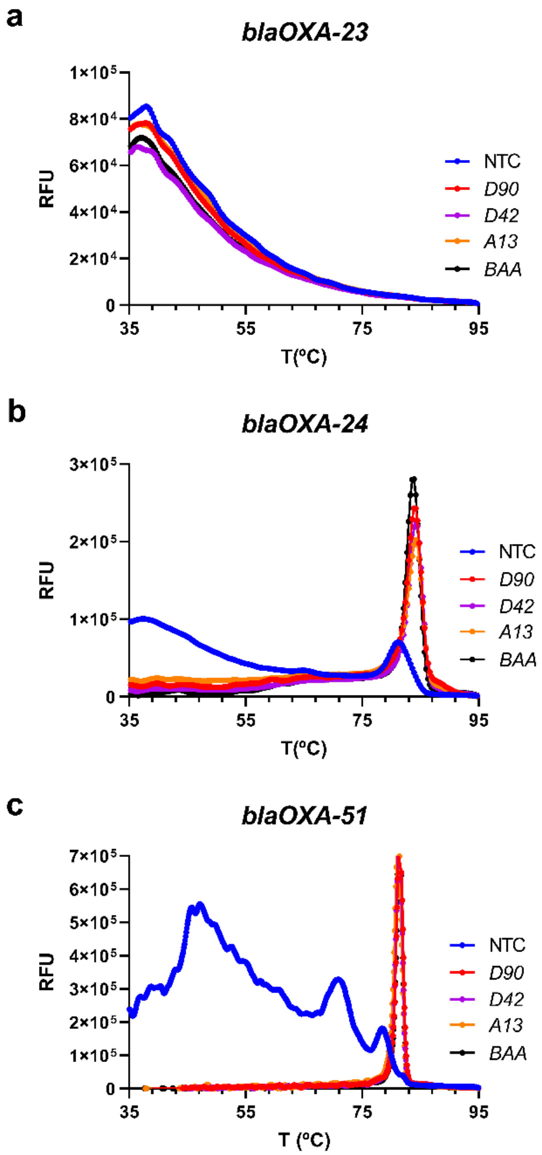


**Supplementary Figure 7** | Melting curve analysis from the LAMP reactions of clinical samples derived from *A. baumannii*. Melting curve for gene *blaOXA-23* (a), *blaOXA-24* (b) and *blaOXA-51* (c) using the primers sets for the 3 different genes. NTC – Non-template-control (negative); D90, D42, A13 are Japanese clinical isolates whereas BAA is BAA-1794 MDR reference strain. LAMP reactions were performed at 65 °C for 1 h, with fluorescence recorded every 1 min. Melt curve analysis was performed from 35 to 95 °C with a heating ramp of 0.3 °C/s. Data represent mean ± s.d. from at least three independent replicates.

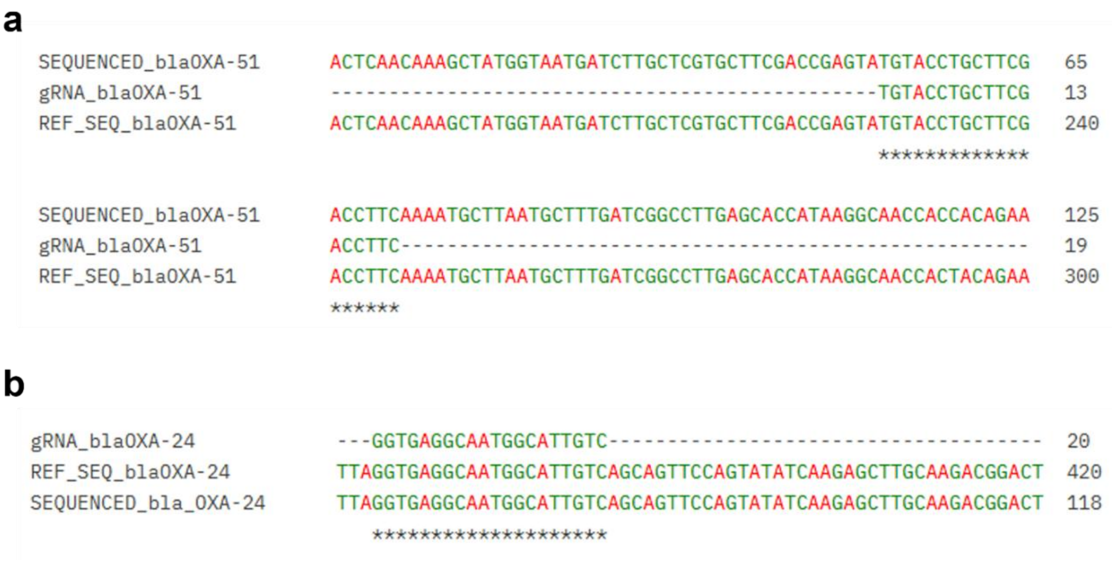


**Supplementary Figure 8 | Conservation of guide RNA target sites in blaOXA genes across clinical *A. baumannii* isolates.** (a) Multiple sequence alignment of the guide RNA protospacer region within the *blaOXA-24* gene from clinical *A. baumannii* genomic DNA. The reference sequence (REF_SEQ_blaOXA-51), Sanger-sequenced amplicons (SEQUENCED_blaOXA-51), and the corresponding guide RNA sequence (gRNA_OXA-51) are shown. (b) Multiple sequence alignment of the guide RNA protospacer region within the *blaOXA-24* gene, presented as in panel A. Sequences were aligned using the The EMBL-EBI Job Dispatcher sequence analysis tools framework (https://www.ebi.ac.uk/jdispatcher)[48]
